# Human insula neurons respond to simple sounds during passive listening

**DOI:** 10.1101/2025.03.12.642819

**Authors:** Joel I Berger, Hiroto Kawasaki, Matthew I Banks, Sukhbinder Kumar, Matthew A Howard, Kirill V Nourski

## Abstract

The insula is critical for integrating sensory information from the body with that arising from the environment. Although previous studies suggest that posterior insula is sensitive to sounds, auditory response properties of insula neurons have not previously been reported. Here, we provide the first report of a population of human single neuron data from the insula and provide comparative data from the primary auditory cortex, recorded intracranially from human participants during passive listening. In each condition, more than 330 single neurons were recorded in 11 participants. Almost a third of neurons in posterior insula and a smaller subset in anterior insula responded to simple tones and clicks. Responsive neurons were distributed throughout posterior and anterior insula and showed preferred frequency tuning. Onset latencies in the insula were similar to those in the primary auditory cortex but response durations were significantly shorter. Overall, these data highlight that insula neurons respond to auditory stimuli even in non-behaviorally relevant contexts and suggest an important contribution of audition to the postulated integrative functions of insular cortex.

## Introduction

The insula, buried within the lateral fissure, is a key brain region involved in processing of interoceptive signals [1-5] which includes sensations not only from within the interior or viscera of body (e.g. cardiac, respiration, hunger) but also those arising from the skin (e.g. pain, temperature and affective touch; [1, 6] for a review, see [7]). Theoretical models of insula function suggest that these interceptive sensations form the basis for subjective feelings [8] and sense of bodily self [9]. By contrast, empirical evidence for the role of insula in processing of exteroceptive signals, such as sounds that are external to the body and not associated with interoceptive signals, remains scant, although there is some limited evidence implicating the insula in auditory processing. Case reports have shown that damage to the insula region can cause alterations in auditory perception [10-13], even without apparent damage to auditory cortex. Additionally, studies in animals [14-18], human neuroimaging using functional magnetic resonance imaging (fMRI) [19-22] and electrophysiology [23-29], suggest that activity within the insula may be modulated by sound, particularly when it is emotionally arousing (e.g. music) or behaviorally relevant (e.g. detection of an odd tone), while recent intracranial local field potential data highlight that posterior insula represents phonological features during both speech production and perception [30]. In misophonia, a condition characterized as an intense emotional reaction to sounds of a particular nature (e.g. chewing), the anterior insula is hyperactivated and hyperconnected when listening to trigger sounds [31]. This finding has been interpreted within the context of social cognition and interoceptive awareness [32, 33]. The insula has also been implicated in auditory hallucinations [34], and intracranial stimulation of this region can produce auditory sensations [35-37], although not always [38]. However, electrophysiological and imaging data have almost always been obtained in behaviorally salient contexts, such as music or speech, while lesion studies may not always be specifically localized to insula exclusively, and there are currently no reports of single neuron responses recorded in the human insula.

Prior studies also do not address whether there is any tuning to basic acoustic sound attributes in the insula [39]. For example, neurons in primary auditory cortex are tuned to specific spectral and temporal features of acoustic stimuli [40-42]. Can activity of individual neurons in the insula also be modulated according to these fundamental acoustic properties even when stimuli are not behaviorally relevant? If so, what are the latencies and tuning properties of these neurons, and where are these located?

We addressed this gap in knowledge by recording extracellular single neuron activity from the insula in eleven humans and comparing these recordings to those obtained from the adjacent posteromedial portion of Heschl’s gyrus (HGPM), corresponding to primary auditory cortex [43]. Data were collected while participants passively listened to auditory stimuli consisting of short click trains and pure tones of various frequencies.

This allowed us to determine (1) the extent of insula neurons that are responsive to auditory stimulation; (2) whether these neurons showed frequency tuning; (3) the spatial distribution of responsive neurons across posterior and anterior insula (InsP and InsA, respectively); and (4) the latencies, magnitudes, and durations of these responses compared to HGPM. Knowledge about the neuronal properties of this brain area in response in audition is important for understanding how the human brain processes sounds. This knowledge has additional relevance to auditory hallucinations and disorders such as tinnitus [44].

## Results

Electrodes were implanted for clinical purposes of monitoring seizures and were localized based on pre- and post-operative neuroimaging (see *Methods*). Single neuron responses to 40 ms 100 Hz click trains were recorded in a single 100-trial block.

Responses to 300 ms tones were recorded in a separate block while participants were presented with pseudo-randomly ordered tones of varying frequency, from 0.25 kHz to 8 kHz, separated by octave steps. Each tone was presented 50 times, resulting in 300 trials total. Interstimulus intervals were 2 s in both experiments (see *Methods* for full details of auditory stimulus presentation).

**Figure 1a** displays the locations from which single neurons were successfully isolated, shown on a template reconstruction of the insula. The size of each marker indicates the number of neurons isolated at a particular location. For click stimuli, a total of 331 neurons were isolated (57 HGPM neurons, 161 InsP neurons and 113 neurons in InsA). For pure tone stimuli, a total of 339 neurons were isolated (59 HGPM neurons, 156 InsP neurons and 124 InsA neurons). **Figure 1b** shows spike waveforms, trial rasters, and kernel-smoothed spike density functions for exemplar neurons in HGPM, InsP and InsA (top to bottom panels, respectively) in response to click train stimuli. In these examples, HGPM shows a sharp onset response to the click train, while the InsP neuron responds consistently across trials with a slightly longer latency than HGPM, and the InsA neuron responds with an even longer latency, with visibly less consistent timing across trials.

**Figure 1:**
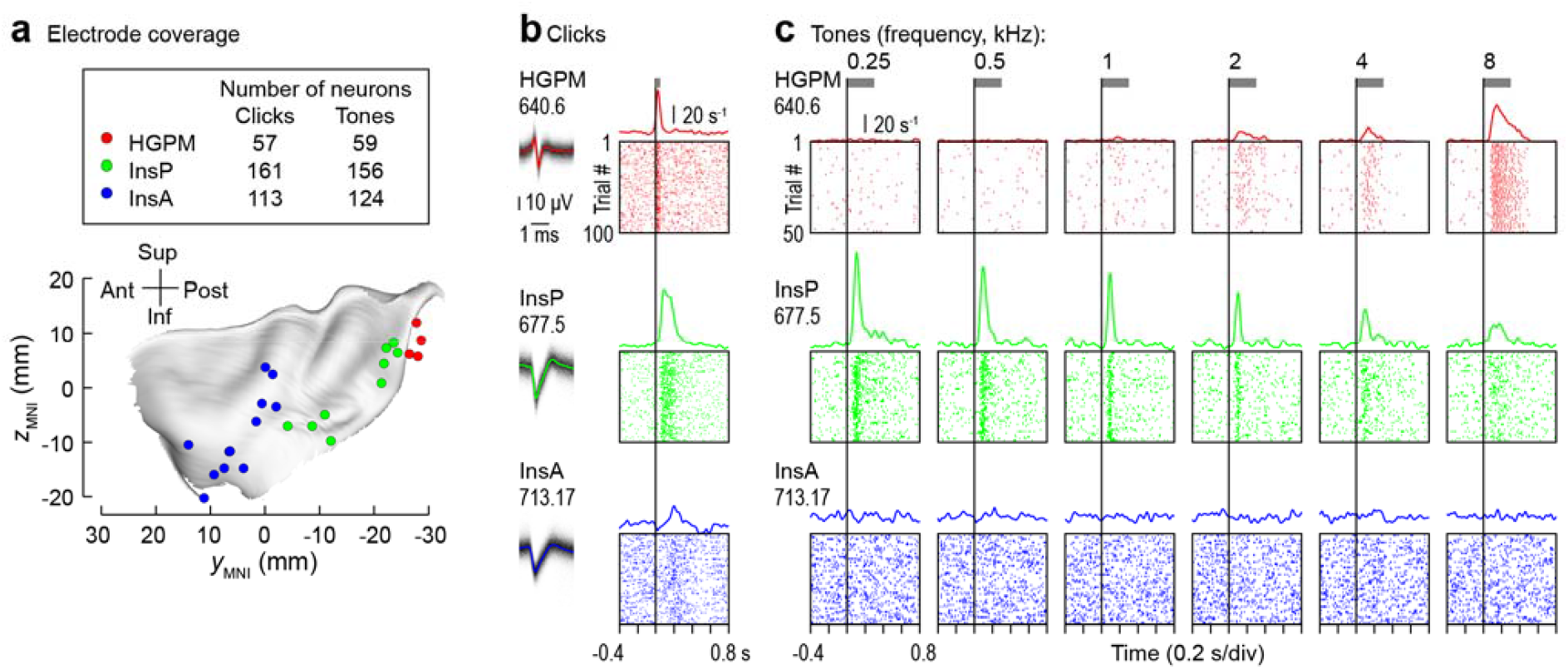
Insula neurons respond to clicks and tones. **a**, Locations and numbers of isolated putative single neurons in response to clicks and tones, plotted on a template reconstruction of insula. **b**, Example responsive neurons in HGPM (top panel), InsP (middle panel) and InsA (bottom panel). For each example, corresponding spike density functions are shown above raster plots. Codes under location names indicate neuron ID. Grey horizontal bars indicate duration of the stimulus. Time on x-axis is relative to stimulus onset (vertical line). **c**, Example responsive neurons for the same locations in response to tones of differing frequencies. For these examples, the corresponding best frequencies were 8 kHz, 0.25 kHz and 2 kHz, respectively, with the InsA neuron exhibiting significant suppression to the tonal stimulus. Throughout, red: HGPM, green: InsP, blue: InsA.

**Figure 1c** shows examples for three single neurons in these same brain locations in response to pure tone stimuli of different frequencies. The neurons in HGPM and InsP showed clear evidence of frequency tuning (to 8 kHz and 0.25 kHz, respectively). Tuning is less obvious in the InsA neuron, though significant suppression was observed in response to tonal stimuli of 2 kHz.

Information such as response latencies can provide crucial information about the position of a region within the hierarchy of an information processing network [45-48]. Thus, to characterize the properties of neuronal responses to clicks across the population, we examined response magnitudes, latencies, and durations for modulated neurons in each region. For each neuron, significant modulation in response to auditory stimuli was determined by comparing the spike rates following the stimulus to the pre-stimulus baseline (see Methods). Trial-averaged raster plots for all significantly modulated neurons in each of the three regions are shown in **Figure 2a**. These are separated according to whether they increased or decreased their activity (indicated by the arrow directions). The overall proportion of modulation response types is then shown in **Figure 2b**. As is evident from both plots, the majority of modulated neurons increased their firing rates in response to click-train stimuli. Across all recorded neurons, 30.4% of neurons in InsP were click-responsive (26.7% increasing, 3.73% decreasing) and 15.9% were modulated to clicks in InsA (9.73% increasing, 6.19% decreasing). By comparison, 84.2% of neurons in HGPM modulated significantly to clicks (77.2% increasing their activity, 7.02% decreasing). Fisher’s Exact probability tests with the Freeman–Halton extension (to allow for preserving modulation direction with a 2 × 3 test) revealed that there were significant differences in the proportions of modulated neurons between HGPM and both InsP and InsA (*p* < 0.0001 for both comparisons), and between InsP and InsA (*p* = 0.001), highlighting that a significantly larger proportion of neurons were modulated in InsP than in InsA, while both regions were proportionally less modulated than HGPM.

**Figure 2:**
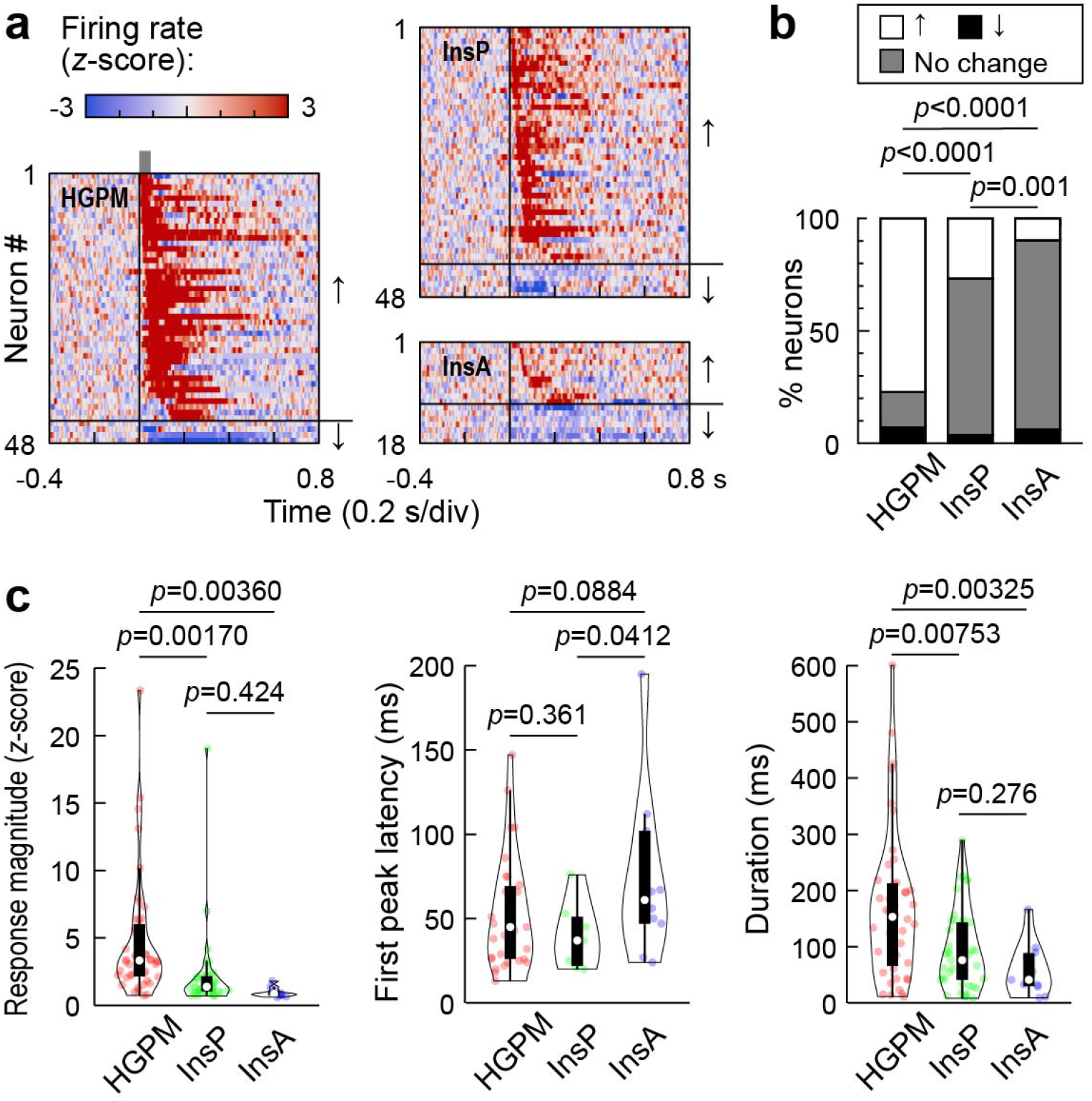
Insula responses to clicks are prominent and have similar latencies to HGPM. **a**, Averaged raster plots for all modulated neurons, organized by region and sorted according to first peak latency. Arrows indicate neurons that either increased (up arrow) or decreased (down arrow) in response to clicks. **b**, Proportion of response types to click-trains for the three different regions. *P*-values based on Fisher’s Exact probability tests. **c**, Violin plots showing response magnitudes (left panel), latencies (middle panel) and durations (right panel) for all three regions, across positively modulated neurons. White symbols denote medians, bars indicate Q_1_ and Q_3_, and whiskers show the range of lower and higher adjacent values (i.e., values within 1.5 interquartile ranges below Q_1_ or above Q_3_, respectively). *P*-values based on linear mixed effects models.

The magnitudes, latencies and durations of all positively modulated responses to click-train stimuli are shown in **Figure 2c**. The average magnitudes of responses following stimulus presentation (0 to 350 ms post-stimulus onset) in InsP and InsA were significantly smaller than HGPM (*p* = 0.00170 and *p* = 0.00360, respectively), with no significant difference between InsP and InsA (*p* = 0.424). Positively modulated InsP neurons exhibited similar latencies to HGPM neurons (*p* = 0.361), with median latencies of 44.0 ms (interquartile range [IQR] = 48.8) and 48.5ms (IQR = 45.5), respectively (middle panel of 2C), which is suggestive of parallel hierarchical auditory processing between the two regions, while InsA latencies were notably slower (median = 67.0 ms, IQR = 57.2). InsP and InsA responses were significantly more transient than HGPM responses (*p* = 0.008 and *p* = 0.003, respectively; right panel in **Figure 2c**), with InsP showing a median duration of 76 ms (IQR = 102.0) compared to a median of 153 ms (IQR = 146.5) in HGPM, highlighting that there were differences in response patterns between insula and HGPM neurons. InsA responses were even more transient (median = 41 ms, IQR = 58.5), though there was no significant difference overall between the response durations of InsP and InsA neurons (*p* = 0.276). Modulated neurons were distributed throughout each subregion of insula, though there was an absence of modulation in the most ventral aspects of InsA (see **Supplementary Figure 2**).

Figure 3 shows the response properties of the different regions for tones. Approximately one third of InsP were modulated by tones (30.8% total; 28.8% increasing, 1.92% decreasing). A higher proportion of neurons in HGPM were modulated by tones (94.9% total; 84.8% increasing, 10.2% decreasing), with a lower proportion in InsA modulated in response to tones compared to clicks (8.07% total; 5.65% increasing, 2.42% decreasing). As with click trains, there were significantly higher proportions of modulated neurons in HGPM compared to both InsP and InsA (*p* < 0.0001 for both comparisons) and in InsP compared to InsA (*p* < 0.0001). See **Supplementary Figure 3** for the spatial distribution of the proportion of tone-modulated neurons.

**Figure 3:**
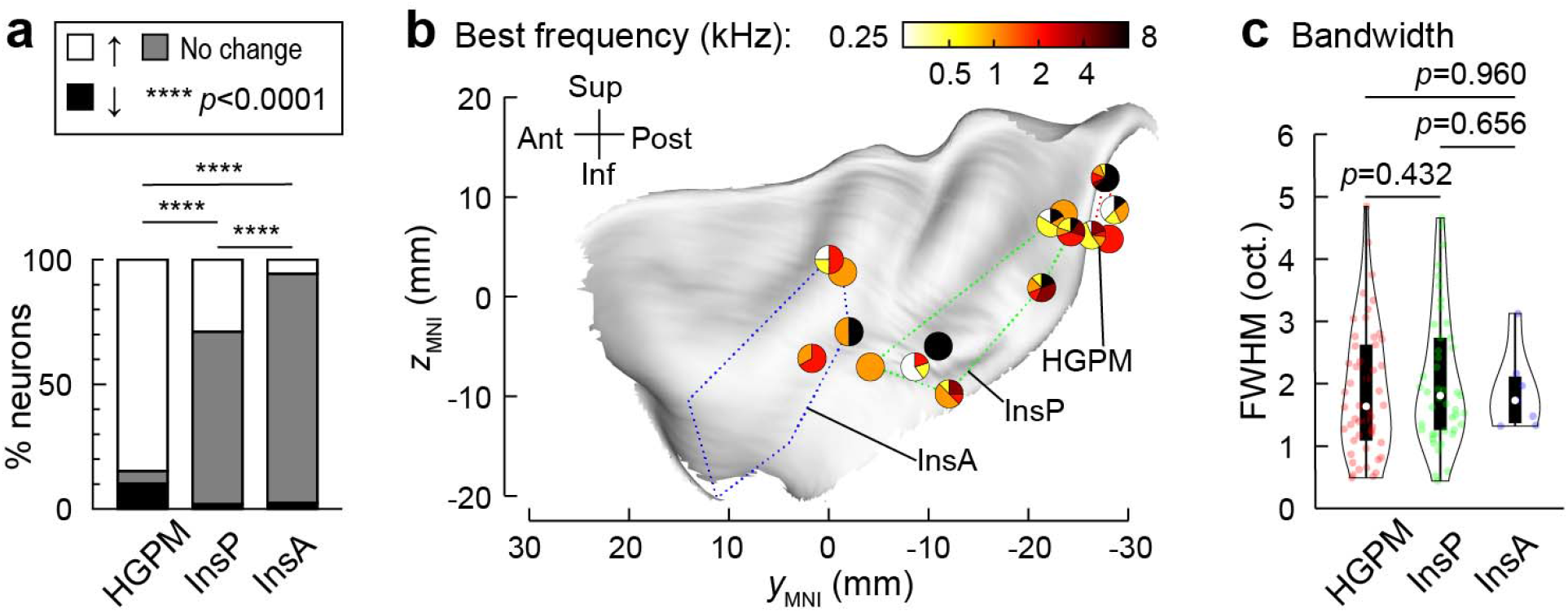
Tuning properties of insula and HGPM neurons. **a**, Proportion of response types for the three different regions. *P*-values based on Fisher’s Exact probability tests. **b**, Proportion of modulated neurons showing best responses to each tone frequency for each recording location. Each marker represents a recording location, with the proportion of modulated neurons responding to different frequencies represented according to the pie chart within that marker, i.e. if 100% of modulated neurons responded to 1 kHz, this would be represented by a fully orange circle. **c**, Violin plots showing bandwidth of all modulated neurons, measured in octaves according to full-width half maximum (FWHM; see Methods). White symbols denote medians, bars indicate Q1 and Q3, and whiskers show the range of lower and higher adjacent values (i.e., values within 1.5 interquartile ranges below Q1 or above Q3, respectively). *P*-values based on a linear mixed effects model.

We examined whether there were differences in frequency tuning for the different regions by determining the proportion of neurons showing a preference to each of the tone frequencies (**Figure 3b**). As the electrodes used in this study do not allow for determining the precise spatial location of wires relative to one another (they only show as a cluster of microwires on the magnetic resonance images [MRI] and do not clearly map to particular channels), along with the relative sparsity of sampling of certain areas of insula, we could not reliably obtain tonotopic maps for the individual electrode arrays. However, as with clicks, there was an absence of response modulation in ventral anterior insula. There was no clear preference for a particular tone frequency across neurons, although fewer neurons in HGPM modulated maximally to 4 kHz than other frequencies. To determine whether there were differences in how precisely tuned insula neurons were, we assessed the bandwidth of responses to different tones (see Methods). This is shown in **Figure 3c** for the three different regions, with the spatial distribution shown in **Supplementary Figure 3**. The median bandwidth for positively modulated neurons was 1.64 octaves for HGPM (IQR = 1.54), 1.81 octaves for InsP (IQR = 1.48) and 1.73 octaves for InsA (IQR = 0.75). There was no significant difference in bandwidth between InsP and InsA (*p* = 0.656), nor between either region and HGPM (*p* = 0.432 and 0.960, respectively), highlighting that the tuning properties to tones were similar across the three regions.

## Discussion

Our results demonstrate that neurons in the human insula significantly modulate their activity in response to auditory stimuli that are passively presented, in the absence of any obvious behavioral or emotional salience. Responses in insula had similar latencies and bandwidths to HGPM, but were more transient. The highest proportion of these responses were in InsP, though a smaller minority of neurons in InsA were also modulated, albeit more weakly. Overall, these results suggest that processing of basic properties of acoustic stimuli contribute to the postulated role of the insula.

The insula, particularly InsA, is known to be part of a salience network [49, 50] which detects and marks how behaviorally relevant the stimulus/events in the environment are. Depending on the degree of saliency, the insula mediates further processing by directing attention or allocating working memory by coordinating between different brain networks (e.g. activating a fronto-parietal network for attention and deactivating the ‘mind wandering’ default mode network for a highly salient event). How does InsA receive sensory inputs for marking their saliency? Theoretical models of insula function [1, 50, 51] suggest that InsP receives sensory inputs which are then re-represented in InsA. There is now ample evidence to support the idea that interoceptive signals from the body do indeed reach InsP [1]. Whether this is also the case for exteroceptive signals such as auditory, however, has historically been less certain. Our high spatial-temporal resolution single neuron data recorded from the insula highlights that – like signals from the body – auditory signals from the external environment are received in InsP. It should be noted that in the saliency model of insula function, stimuli need not be salient or behaviorally relevant to reach InsP, and they were not in the case of the presence study. The saliency of the stimuli is proposed to be marked in InsA. This model may then explain why in our data the proportion of modulated neurons is less in InsA compared to that in InsP when passively listening to sounds, as these sounds were not required to be salient.

Evidence showing that insula activity is modulated even during passive listening may explain – at least in part – why lesions to the insula can cause altered auditory processing, such as in cases of acquired amusia, temporal processing deficits or hyperacusis after a stroke [52, 53]. The results of the current study are also consistent with neuroanatomical studies showing that the insula receives direct projections from auditory cortex [54, 55] and medial geniculate body [56], although as highlighted by Remedios et al. [14], anatomical results do not give insight into the nature of the involvement of the insula in auditory processing without corresponding physiological data, which we have provided here.

These results may also explain why stimulation of insula can produce auditory perceptual experiences [35-37]. Contrasting with these other studies, Duong et al. [38] did not find auditory responses from stimulation of the insula, which the authors suggested was due to previous studies stimulating structures adjacent to rather than within the insula proper. In the current study, our electrode locations were confirmed to be in the insula based on individual participants’ MRIs, as well as plotted on a template using MNI coordinates. While it is plausible that some studies may have stimulated regions adjacent to the insula to produce auditory perceptual experiences, another possibility is that an absence of behavioral response may be due to the still relatively small fraction of neurons in this region being involved in auditory processing (about 1/3 in the current study). Thus, if particular neurons were not stimulated by electrodes within the insula, then an auditory percept would likely not be produced. Further studies utilizing both stimulation mapping and auditory presentation in the same neurons would be invaluable for elucidating the dynamics of these auditory-responsive insula neurons.

Reports of the tuning properties of single neurons in HGPM are rare, with only a handful studies to date examining this [57-59]. In the first systematic study of individual neurons in human auditory cortex, the bandwidth measured in a similar manner to the current manuscript was at least 1 octave [58], consistent with what we found here. Contrastingly though, the bandwidth of neuronal responses to tones here was considerably larger than what was demonstrated previously when using an approach that estimates based on spectro-temporal receptive fields derived from random-chord stimuli [59]. It is therefore likely that basic tonal stimuli – although widely used historically to study the function of the auditory system [60] – do not reveal the full extent of the precision of stimulus encoding, and future studies should examine this with refined stimuli for both the HGPM and the insula. Moreover, while this study represents a rich and unique dataset, the spatial extent of electrode coverage was still somewhat limited, as determined solely by the clinical needs of the patients. This work lays the foundation for further, more comprehensive, studies aimed at understanding the full extent of auditory-related single neuron activation throughout the insula.

## Methods

### Participants

Extracellular single neuron recordings were obtained from 11 adult neurosurgical patients, implanted with electrodes for the purposes of monitoring epileptiform activity prior to potential treatment. Research was conducted under approval of the University of Iowa Institutional Review Board and written informed consent was obtained from all participants prior to data collection. Recordings were made while subjects were reclined in a hospital bed, in a custom-designed dedicated electromagnetically-shielded facility within the University of Iowa Clinical Research Unit.

### Electrodes and recording system

Stereo electroencephalography (sEEG) depth electrodes (Ad-Tech Medical, Oak Creek, WI) were placed in brain locations based solely on a clinical need to identify seizure foci [61]. For the purposes of recording single neurons, sEEG electrodes included here in the insula and HGPM were of a hybrid design [62]. These consisted of eight 39 µm diameter platinum-iridium high-impedance microwires that were insulated plus one uninsulated microwire. These microwires protruded from the end of the macro recording probe and were prepared with a cut length between 2 to 4 mm, depending on the distance of the most distal macro contact to the appropriate brain target. Each of these microwires was individually separated in a splay pattern in the operating room immediately prior to implantation. Electrode locations were confirmed based on post-operative MRI scans, preprocessed using Freesurfer [63] (see below for further details). All neurophysiological data were recorded using a Neuralynx Atlas System (Neuralynx, Bozeman, MT). High impedance recordings were first passed through a preamplifier located on top of the patient’s head (ATLAS-HS-36-CHET-A9, Neuralynx, Bozeman, MT) prior to interfacing with the ATLAS acquisition system. These were subsequently recorded with a 32000 Hz sampling rate, filtered between 0.1 – 8000 Hz and referenced online to the uninsulated microwire.

### Stimuli and procedure

Experimental stimuli were trains of acoustic clicks (used previously in [64]) and pure tones (used previously in [65]). Clicks were digitally generated as rectangular pulses (0.2 ms duration) and were presented in trains of five at a rate of 100 Hz (train duration 40 ms, 2 s inter-train interval). Tones were presented at six frequencies between 250 and 8 kHz in 1-octave steps (300 ms duration, 5 ms rise–fall time, 2 s interstimulus interval). The six tones were presented 50 times each in a random order. The stimuli were presented at a comfortable level, approximately 50-60 dB above hearing threshold. Stimuli were delivered to both ears via insert earphones (ER4B, Etymotic Research, Elk Grove Village, IL) that were integrated into custom-fit earmolds or foam ear tips (for two participants). In each participant, the intensity difference between click trains and pure tones was within 20 dB and varied depending on comfort. Inter-stimulus intervals were chosen randomly within a Gaussian distribution (mean interval 2 s; SD = 10 ms) to reduce heterodyning in the recordings secondary to power line noise. Stimulus delivery was controlled by a TDT RP2.1 and RZ2 real-time processor (Tucker Davis Technologies, Alachua, FL).

### Imaging

A T1-weighted structural MRI scan of the brain was conducted for each participant both before and after the implantation of electrodes. The images were captured using a 3T Siemens TIM Trio scanner and a 12-channel head coil. MPRAGE images had a spatial resolution of 0.78 × 0.78 mm, a slice thickness of 1.0 mm, and utilized a repetition time (TR) of 2.53 s and an echo time (TE) of 3.52 ms. To locate the recording positions on the preoperative structural MRI scans, these images were aligned with post-implantation structural MRIs. This alignment was achieved using a 3D linear registration algorithm (Functional MRI of the Brain Linear Image Registration Tool; [66]) and custom-written MATLAB scripts (MathWorks, Natick, MA). Included microwire bundles were verified to be within gray matter for either the insula or HGPM. Electrode locations were co-registered for each participant to a template brain, in order to derive MNI coordinates for the purposes of visualization. Coordinates of microwire locations were then visualized on an fsaverage MRI template showing the insula and neighboring region of HGPM, using custom-written MATLAB scripts.

### Data processing and analysis

Data from high impedance electrodes localized to the insula or HGPM were first extracted using Matlab and denoised with an implementation of the demodulated band transform [67]. These data were downsampled to 12 kHz and common average re-referenced to all high impedance contacts on the same assembly prior to spike sorting. Spike sorting was performed using an automated procedure with manual curation, utilizing an algorithm implementing high-order spectral decomposition to aid in pattern recognition and identify individual features [68]. Briefly, filters were estimated for potential candidate single neuron waveforms for each channel through a process related to blind deconvolution. Extracted features were clustered using a gaussian mixture model in Matlab (R2022a, Mathworks Inc) and spike times from these clustered features were used to plot separate candidate waveforms. Single neurons were then manually curated and defined based on classical waveform shapes, uniformity of waveforms across different spike times for each cluster, and interspike interval distributions that did not violate refractory periods (<1% of interspike intervals occurring within 1 ms). See **Supplementary Figure 1** for summarized curation details. Putative single neuron spike times were then epoched around the stimuli for each trial (500 ms before stimulus onset to 1000 ms post-stimulus onset). Raster plots were created to show neuronal activity for each trial and spike density functions were estimated by convolving single neuron spike times with a gaussian kernel (1 ms resolution, generally 5 ms standard deviation, though a standard deviation of 15 ms was used for display purposes in **Figure 1**).

### Statistical analysis

Single neuron modulation to auditory stimuli was determined using a two-tailed paired *t*-test against baseline, wherein spike rates across trials between 0 to 350 ms post-stimulus were compared to baseline firing rates from a pre-stimulus window of the same duration (−400 to -50 ms). This was only considered for neurons that had a mean firing rate ≥1 Hz across the baseline and post-stimulus windows. The *p*-value alpha for this test was set at 0.05 for click stimuli and Benjamini-Hochberg false discovery rate (FDR) correction [69, 70] was applied to tonal response *p*-values to account for multiple comparisons (based on the six different tone frequencies presented). Magnitudes for all responses were determined by averaging the z-scored firing rate across the post-stimulus window. The latency of significantly modulated neurons was then determined according to the first peak in the baseline z-scored spike density function from 10 ms to 350 ms post-stimulus, wherein this peak was a minimum value of half of the maximum peak within this time window. For neurons that suppressed their activity in response to auditory stimuli, spike density functions were first inverted prior to finding the peak. For tone stimuli, the best frequency of each neuron was defined as the tone that elicited the maximum absolute value of the z-scored average, wherein this response was significant. Bandwidth in octaves was determined based on the full width at half maximum of this frequency, which was obtained after interpolation of both frequencies and magnitudes to create 10000 discrete sampling points. Where the lowest or highest frequencies were the maximum absolute value (i.e. showed the strongest modulation), these were taken as the low or high cutoff for this calculation, respectively. Response durations were determined based on the time after the first peak wherein the spike density function was reduced to 5% of the maximum peak value. Regional differences in response properties were examined using a linear mixed effects modeling approach.

Output (response magnitude, latency or duration) was modeled using region of interest (ROI, i.e. HGPM, InsP, or InsA) as a categorical fixed effect, with random intercept of channel nested within participant:

Output ∼ ROI + (1 | Participant : Channel) + (1 | Participant)

## Data and code availability

Data and analysis scripts used in this manuscript are available upon reasonable request from the authors.

## Acknowledgements

This work was supported by the National Institutes of Health (R01 DC004290, awarded to M.A.H.). We are grateful to Haiming Chen, Phillip E Gander, Christopher M Garcia, Christopher K Kovach and Ariane E Rhone for help with electrode preparation, data collection and design of the spike sorting methodology.

---

**Supplementary Figure 1.**
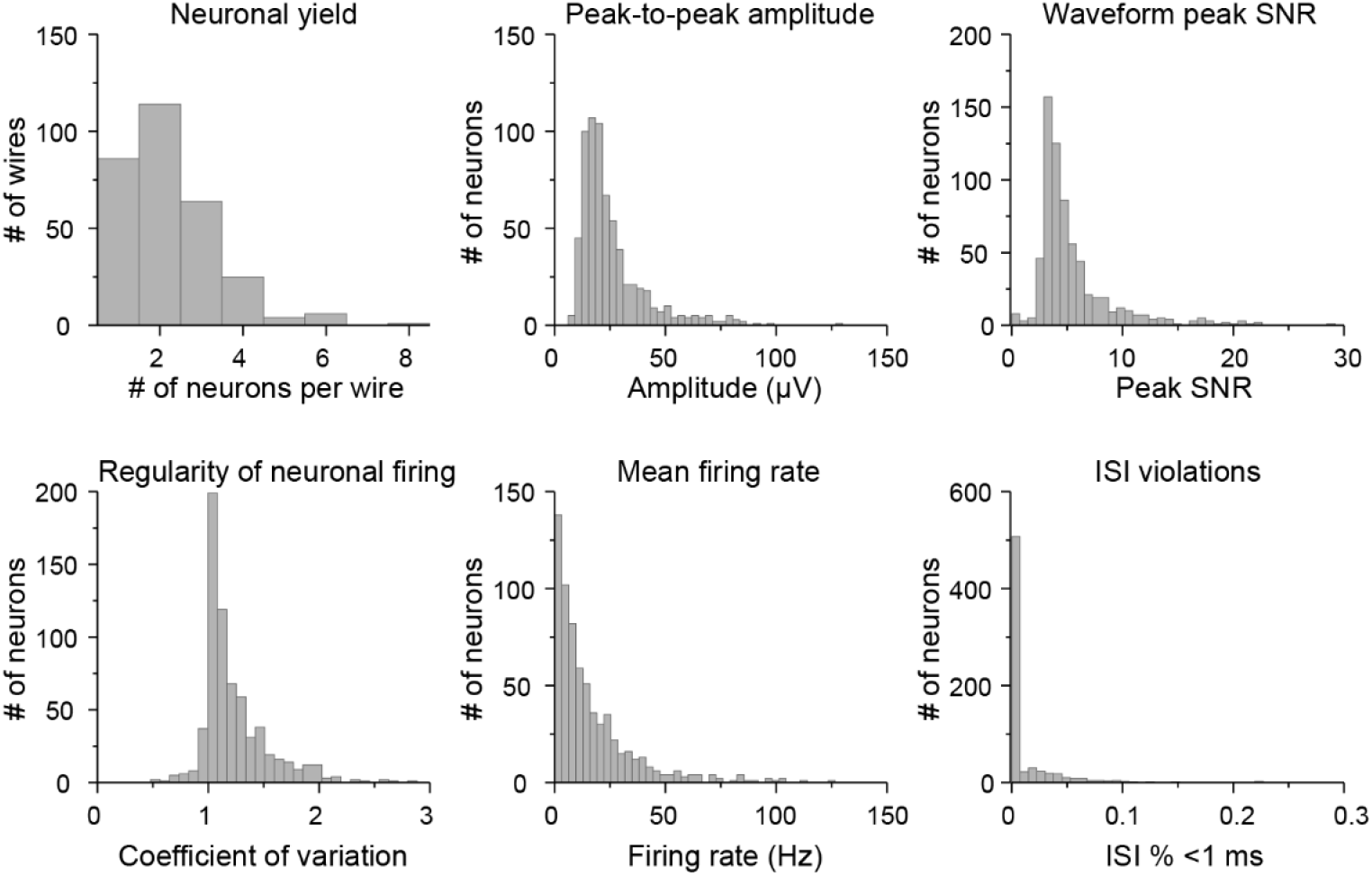
Curation plots for all isolated putatitive single neurons. ISI: inter-spike interval. SNR: signal-to-noise ratio, calculated as the ratio of the peak of each single neuron waveform divided by the median absolute value of the signal for that channel, based on the method implemented in SpikeForest [71].

**Supplementary Figure 2.**
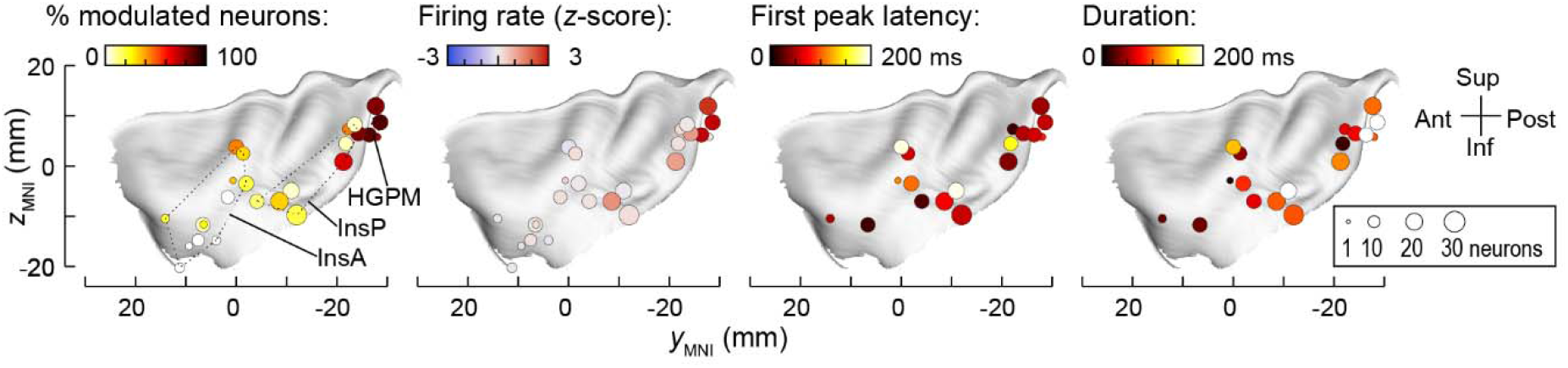
Template reconstruction of insula showing for clicks (left to right) the proportion of modulated neurons, mean magnitude of firing rate changes, latency of modulation and duration of modulation.

**Supplementary Figure 3.**
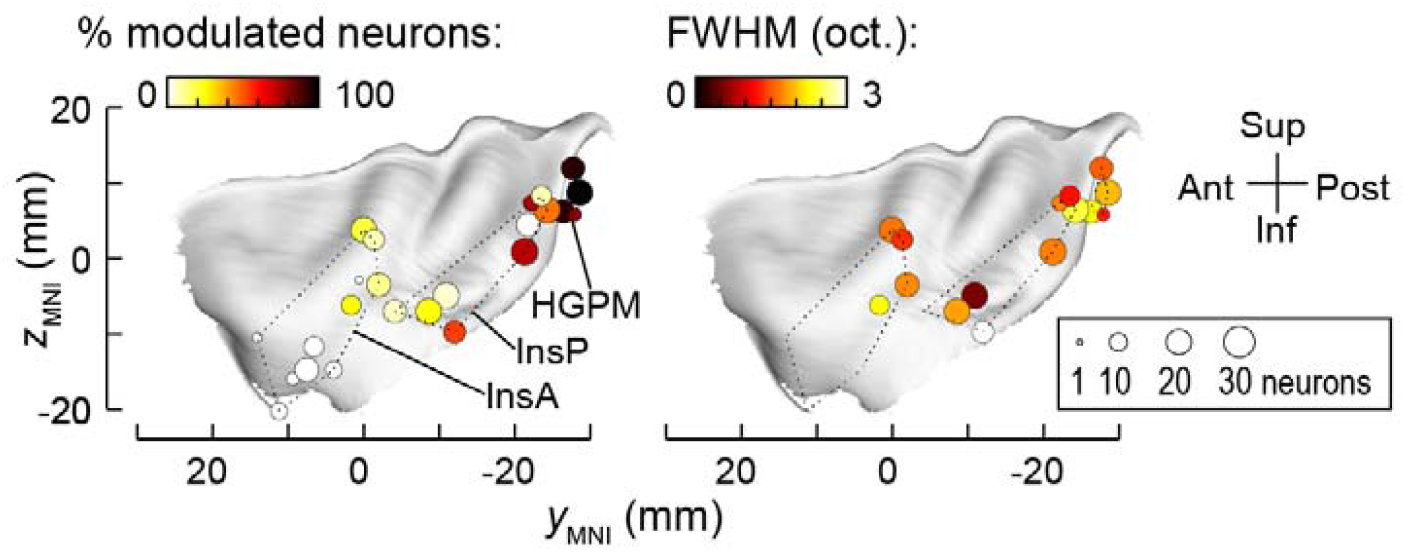
Template reconstruction of insula showing for tones the proportion of modulated neurons (left panel) and mean bandwidth of tuning within a location (right panel).

